## Supplemental Information for "Global patterns of aegyptism without arbovirus"

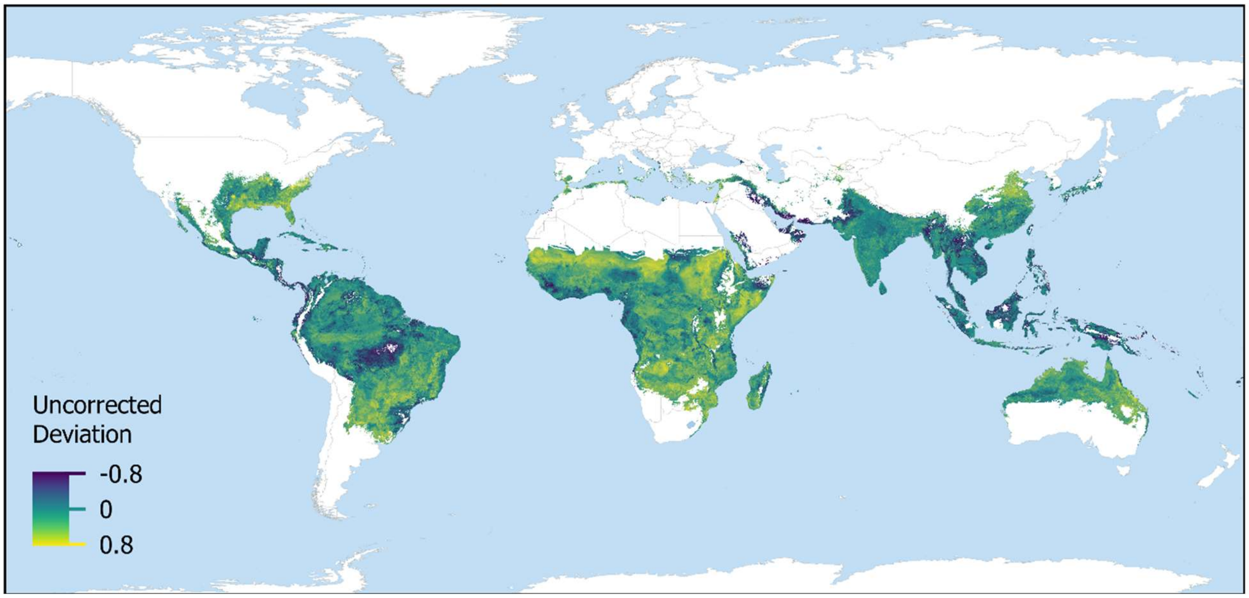


**Supplemental Figure 1.** Uncorrected deviation between *Ae. aegypti* and dengue environmental suitability.

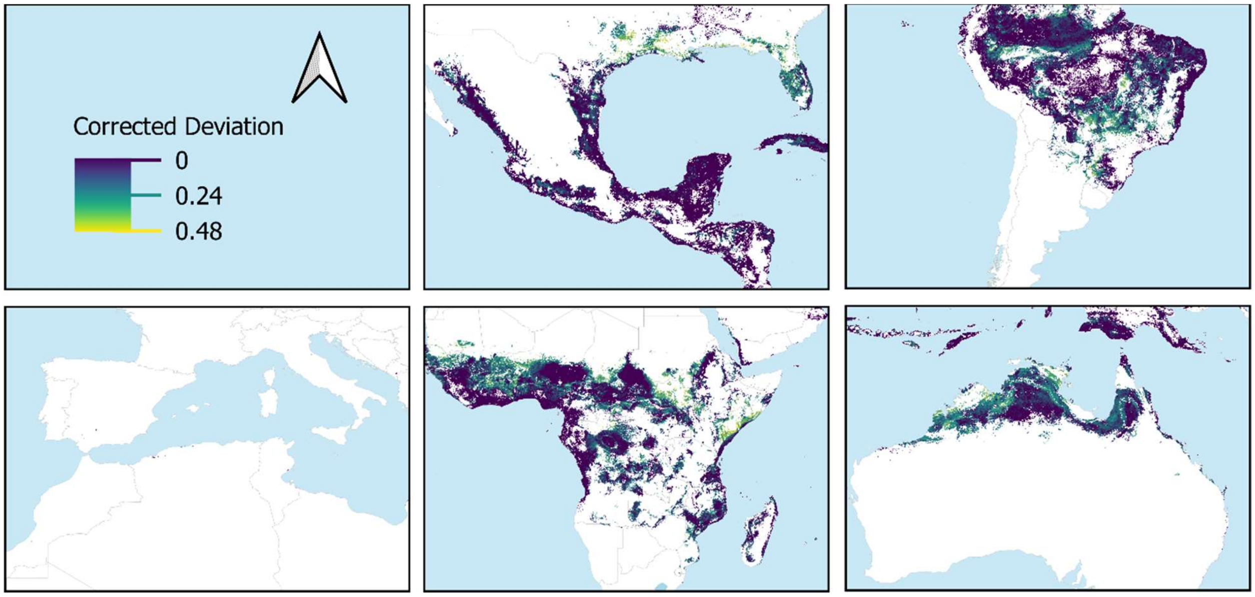


**Supplemental Figure 2.** Deviation between *Ae. aegypti* probability of occurrence and dengue environmental suitability, zoomed in on North America, South America, South Europe and North Africa, Africa, and Australia.

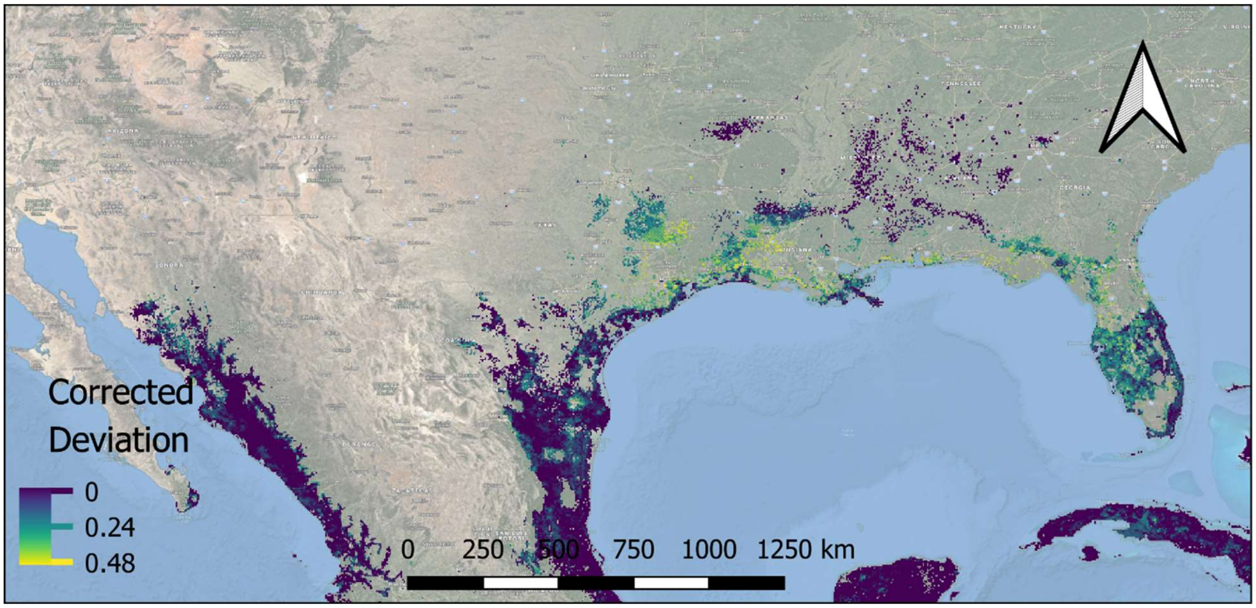


**Supplemental Figure 3.** Deviation between *Ae. aegypti* probability of occurrence and dengue environmental suitability for the Southern United States.

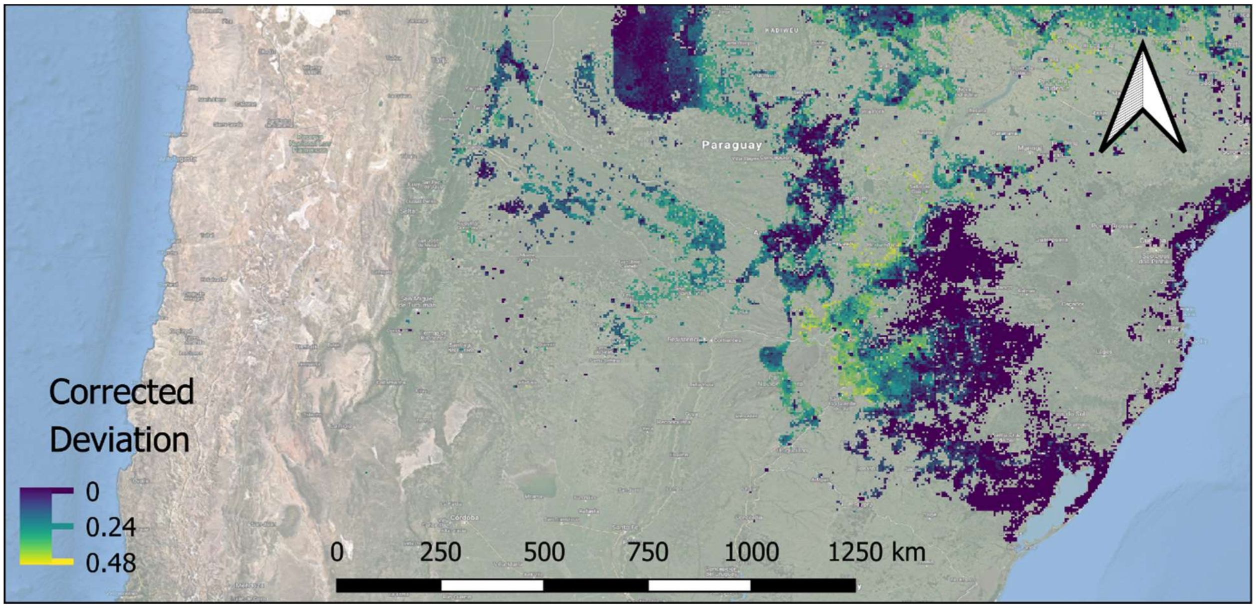


**Supplemental Figure 4.** Deviation between *Ae. aegypti* probability of occurrence and dengue environmental suitability for Northern Argentina, Paraguay, and Southern Brazil.

**Supplemental Table 1.** Data sources for the global rasters used in this paper.

| Variable | Scale | Data Source/URL | Reference |
|---|---|---|---|
| Global distribution of *Ae. aegypti* | 5 x 5 km | https://datadryad.org/resource/doi:10.5061/dryad.47v3c | [1] |
| Global distribution of dengue | 5 x 5 km | https://figshare.com/s/d7d7871d00afe2870619 | [2] |
| Population Density | 1 x 1 km | https://sedac.ciesin.columbia.edu/data/set/gpw-v4-population-density-rev11/data-download | [3] |
| Gross domestic product | 1 x 1 km | https://doi.org/10.5061/dryad.dk1j0 | [4] |
| Infant mortality rate | National /Subnational 1 x 1 km | https://sedac.ciesin.columbia.edu/data/set/povmap-global-subnational-infant-mortality-rates-v2/data-download | [5] |
| Temperature | 1 x 1 km | https://www.worldclim.org/data/worldclim21.html# | [6] |
| Precipitation | 1 x 1 km | https://www.worldclim.org/data/worldclim21.html# | [6] |

**Supplemental Table 2.** Statistical summary of *Ae. aegypti* minus dengue deviation, by country. Blue indicates the lower end of the spectrum, where *Ae. aegypti* occurrence and risk of dengue is nearly equal and high, and yellow represents the other end of the spectrum where *Ae. aegypti* can be found without dengue. Countries with 5 or fewer cells (5 km$^2$) were removed from the table for brevity.

| Country | Count | Mean Deviation | SD | Min | Max | Range |
|---|---|---|---|---|---|---|
| *Mauritania* | 1537 | 0.270822 | 0.076399 | 0 | 0.468075 | 0.468075 |
| *Niger* | 3212 | 0.241343 | 0.075063 | 0 | 0.43299 | 0.43299 |
| *Somalia* | 7144 | 0.210229 | 0.140471 | 0 | 0.476313 | 0.476313 |
| *Burkina Faso* | 9555 | 0.209692 | 0.08242 | 0 | 0.464294 | 0.464294 |
| *eSwatini* | 25 | 0.20527 | 0.030981 | 0.159627 | 0.267192 | 0.107565 |
| *S. Sudan* | 13965 | 0.189314 | 0.126751 | 0 | 0.461959 | 0.461959 |
| *Senegal* | 5230 | 0.182488 | 0.100429 | 0 | 0.449441 | 0.449441 |
| *South Africa* | 93 | 0.182341 | 0.080277 | 0 | 0.359835 | 0.359835 |
| *Botswana* | 15 | 0.182296 | 0.057185 | 0.084908 | 0.296197 | 0.211289 |
| *Kenya* | 8705 | 0.176906 | 0.130131 | 0 | 0.466411 | 0.466411 |
| *Argentina* | 3983 | 0.176614 | 0.105083 | 0 | 0.439078 | 0.439078 |
| *Netherlands* | 9 | 0.176019 | 0.089169 | 0 | 0.273538 | 0.273538 |
| *United States of America* | 12322 | 0.160234 | 0.127157 | 0 | 0.473787 | 0.473787 |
| *Mali* | 12187 | 0.143336 | 0.096475 | 0 | 0.444084 | 0.444084 |
| *Zimbabwe* | 272 | 0.140671 | 0.082867 | 0 | 0.417163 | 0.417163 |
| *Sudan* | 8478 | 0.133645 | 0.124151 | 0 | 0.44961 | 0.44961 |
| *Togo* | 2485 | 0.131907 | 0.111805 | 0 | 0.460853 | 0.460853 |
| *Paraguay* | 7689 | 0.131267 | 0.110785 | 0 | 0.459871 | 0.459871 |
| *Benin* | 5000 | 0.128714 | 0.099668 | 0 | 0.445118 | 0.445118 |
| *Australia* | 42383 | 0.125245 | 0.093226 | 0 | 0.449273 | 0.449273 |
| *Chad* | 12522 | 0.124446 | 0.122251 | 0 | 0.443918 | 0.443918 |
| *Gambia* | 423 | 0.121773 | 0.076891 | 0 | 0.375483 | 0.375483 |
| *Algeria* | 33 | 0.120733 | 0.114471 | 0 | 0.311777 | 0.311777 |
| *Zambia* | 7392 | 0.119709 | 0.0742 | 0 | 0.319709 | 0.319709 |
| *Ghana* | 9599 | 0.115504 | 0.114871 | 0 | 0.455999 | 0.455999 |
| *Singapore* | 21 | 0.114384 | 0.064616 | 0 | 0.214392 | 0.214392 |
| *Namibia* | 351 | 0.110637 | 0.074253 | 0 | 0.284943 | 0.284943 |
| *Mauritius* | 84 | 0.109631 | 0.099659 | 0 | 0.430635 | 0.430635 |
| *Dem. Rep. Congo* | 49377 | 0.102295 | 0.093354 | 0 | 0.441369 | 0.441369 |
| *Saudi Arabia* | 1314 | 0.098834 | 0.100066 | 0 | 0.42865 | 0.42865 |
| *Guinea-Bissau* | 1155 | 0.098167 | 0.08264 | 0 | 0.423113 | 0.423113 |
| *Mozambique* | 18281 | 0.095912 | 0.088501 | 0 | 0.441035 | 0.441035 |
| *Afghanistan* | 29 | 0.093353 | 0.105213 | 0 | 0.322509 | 0.322509 |
| *Uganda* | 4281 | 0.092651 | 0.087806 | 0 | 0.43509 | 0.43509 |
| *Djibouti* | 585 | 0.091013 | 0.092261 | 0 | 0.380496 | 0.380496 |

**Supplemental Table 2, continued**

| Country | Count | Mean Deviation | SD | Min | Max | Range |
|---|---|---|---|---|---|---|
| *Ethiopia* | 14335 | 0.089882 | 0.103374 | 0 | 0.473594 | 0.473594 |
| *Turkey* | 125 | 0.086916 | 0.103747 | 0 | 0.347339 | 0.347339 |
| *Tanzania* | 12375 | 0.086016 | 0.086437 | 0 | 0.401491 | 0.401491 |
| *Brazil* | 251421 | 0.080467 | 0.09552 | 0 | 0.454705 | 0.454705 |
| *Libya* | 10 | 0.076496 | 0.074334 | 0 | 0.208514 | 0.208514 |
| *Angola* | 13627 | 0.075843 | 0.081154 | 0 | 0.322925 | 0.322925 |
| *Central African Rep.* | 23993 | 0.075756 | 0.086161 | 0 | 0.41307 | 0.41307 |
| *Côte d'Ivoire* | 13634 | 0.075138 | 0.101408 | 0 | 0.46961 | 0.46961 |
| *Congo* | 8203 | 0.074303 | 0.080669 | 0 | 0.437049 | 0.437049 |
| *Bolivia* | 20245 | 0.072595 | 0.086744 | 0 | 0.420899 | 0.420899 |
| *Bangladesh* | 6589 | 0.070617 | 0.080114 | 0 | 0.391463 | 0.391463 |
| *Madagascar* | 10491 | 0.068162 | 0.078189 | 0 | 0.420165 | 0.420165 |
| *Spain* | 7 | 0.067902 | 0.086906 | 0 | 0.252066 | 0.252066 |
| *Malawi* | 1112 | 0.066335 | 0.086921 | 0 | 0.360916 | 0.360916 |
| *Guam* | 12 | 0.064371 | 0.060257 | 0 | 0.163893 | 0.163893 |
| *Israel* | 25 | 0.06422 | 0.102963 | 0 | 0.332563 | 0.332563 |
| *Cameroon* | 12896 | 0.062199 | 0.085394 | 0 | 0.421467 | 0.421467 |
| *Hong Kong* | 29 | 0.061137 | 0.077636 | 0 | 0.302333 | 0.302333 |
| *Colombia* | 36374 | 0.060055 | 0.075168 | 0 | 0.424601 | 0.424601 |
| *China* | 25740 | 0.058021 | 0.078603 | 0 | 0.43586 | 0.43586 |
| *Nigeria* | 40458 | 0.056554 | 0.079571 | 0 | 0.443585 | 0.443585 |
| *Uruguay* | 21 | 0.056547 | 0.063163 | 0 | 0.171148 | 0.171148 |
| *Eritrea* | 1507 | 0.056413 | 0.075275 | 0 | 0.437792 | 0.437792 |
| *Aruba* | 6 | 0.055947 | 0.042776 | 0 | 0.140137 | 0.140137 |
| *Haiti* | 1125 | 0.053313 | 0.081108 | 0 | 0.407427 | 0.407427 |
| *Peru* | 28758 | 0.050836 | 0.0784 | 0 | 0.448863 | 0.448863 |
| *Myanmar* | 18541 | 0.049784 | 0.072837 | 0 | 0.431414 | 0.431414 |
| *Bhutan* | 469 | 0.048846 | 0.067289 | 0 | 0.275594 | 0.275594 |
| *Venezuela* | 28011 | 0.047027 | 0.066544 | 0 | 0.426449 | 0.426449 |
| *Brunei* | 254 | 0.046476 | 0.05883 | 0 | 0.31676 | 0.31676 |
| *India* | 131543 | 0.04542 | 0.067103 | 0 | 0.482278 | 0.482278 |
| *Guinea* | 10106 | 0.044463 | 0.071513 | 0 | 0.463709 | 0.463709 |
| *Guyana* | 7224 | 0.043455 | 0.070662 | 0 | 0.408294 | 0.408294 |
| *Nepal* | 2929 | 0.041801 | 0.066457 | 0 | 0.421316 | 0.421316 |
| *Puerto Rico* | 371 | 0.040727 | 0.057873 | 0 | 0.288344 | 0.288344 |
| *Burundi* | 67 | 0.038958 | 0.061973 | 0 | 0.255066 | 0.255066 |
| *Honduras* | 3574 | 0.037542 | 0.071488 | 0 | 0.374007 | 0.374007 |
| *Japan* | 798 | 0.03687 | 0.069031 | 0 | 0.45001 | 0.45001 |
| *Taiwan* | 1023 | 0.036766 | 0.05761 | 0 | 0.458133 | 0.458133 |
| *Mexico* | 30091 | 0.036353 | 0.062812 | 0 | 0.442393 | 0.442393 |

**Supplemental Table 2, continued**

| Country | Count | Mean Deviation | SD | Min | Max | Range |
|---|---|---|---|---|---|---|
| *Liberia* | 2271 | 0.03601 | 0.063775 | 0 | 0.353039 | 0.353039 |
| *Greece* | 38 | 0.035851 | 0.083678 | 0 | 0.328919 | 0.328919 |
| *Cuba* | 5161 | 0.03374 | 0.054247 | 0 | 0.357021 | 0.357021 |
| *Pakistan* | 11821 | 0.033113 | 0.0626 | 0 | 0.435065 | 0.435065 |
| *Cambodia* | 6908 | 0.032395 | 0.057148 | 0 | 0.394668 | 0.394668 |
| *Sierra Leone* | 2137 | 0.032228 | 0.050699 | 0 | 0.309454 | 0.309454 |
| *Suriname* | 3890 | 0.028387 | 0.04457 | 0 | 0.309013 | 0.309013 |
| *Dominican Rep.* | 1649 | 0.027419 | 0.053722 | 0 | 0.341377 | 0.341377 |
| *New Caledonia* | 856 | 0.026595 | 0.045528 | 0 | 0.27335 | 0.27335 |
| *Tonga* | 17 | 0.02625 | 0.043202 | 0 | 0.131397 | 0.131397 |
| *Curaçao* | 20 | 0.023596 | 0.036404 | 0 | 0.111911 | 0.111911 |
| *Barbados* | 18 | 0.023528 | 0.032648 | 0 | 0.102893 | 0.102893 |
| *Thailand* | 20176 | 0.022727 | 0.044623 | 0 | 0.416792 | 0.416792 |
| *Syria* | 10 | 0.02266 | 0.037387 | 0 | 0.109027 | 0.109027 |
| *Yemen* | 1884 | 0.020873 | 0.049575 | 0 | 0.345343 | 0.345343 |
| *Papua New Guinea* | 9466 | 0.020219 | 0.047637 | 0 | 0.414984 | 0.414984 |
| *Gabon* | 7059 | 0.020012 | 0.046014 | 0 | 0.277781 | 0.277781 |
| *Belize* | 817 | 0.019791 | 0.043049 | 0 | 0.295161 | 0.295161 |
| *Laos* | 3776 | 0.018606 | 0.04681 | 0 | 0.302114 | 0.302114 |
| *Turks and Caicos Is.* | 11 | 0.017997 | 0.039329 | 0 | 0.134937 | 0.134937 |
| *Comoros* | 51 | 0.017759 | 0.039172 | 0 | 0.169279 | 0.169279 |
| *Indonesia* | 61849 | 0.017626 | 0.044067 | 0 | 0.458635 | 0.458635 |
| *Philippines* | 8957 | 0.017602 | 0.036593 | 0 | 0.339598 | 0.339598 |
| *Nicaragua* | 2649 | 0.017339 | 0.045104 | 0 | 0.385047 | 0.385047 |
| *Fr. Polynesia* | 41 | 0.017271 | 0.050807 | 0 | 0.27058 | 0.27058 |
| *Cabo Verde* | 86 | 0.016027 | 0.035636 | 0 | 0.155874 | 0.155874 |
| *Antigua and Barb.* | 7 | 0.016003 | 0.0392 | 0 | 0.112023 | 0.112023 |
| *St. Vin. and Gren.* | 13 | 0.015825 | 0.028461 | 0 | 0.100453 | 0.100453 |
| *Sri Lanka* | 2757 | 0.014808 | 0.034071 | 0 | 0.325308 | 0.325308 |
| *Guatemala* | 2846 | 0.013824 | 0.040273 | 0 | 0.345901 | 0.345901 |
| *Iraq* | 1334 | 0.013004 | 0.040605 | 0 | 0.313613 | 0.313613 |
| *U.S. Virgin Is.* | 9 | 0.012696 | 0.024272 | 0 | 0.067725 | 0.067725 |
| *Ecuador* | 4006 | 0.011557 | 0.040101 | 0 | 0.320929 | 0.320929 |
| *El Salvador* | 661 | 0.010152 | 0.0362 | 0 | 0.251234 | 0.251234 |
| *Vietnam* | 11576 | 0.010046 | 0.029572 | 0 | 0.325532 | 0.325532 |
| *Cayman Is.* | 6 | 0.009814 | 0.021944 | 0 | 0.058882 | 0.058882 |
| *Jordan* | 16 | 0.009607 | 0.020189 | 0 | 0.079458 | 0.079458 |
| *Malaysia* | 9712 | 0.009353 | 0.028345 | 0 | 0.352326 | 0.352326 |
| *Dominica* | 26 | 0.008615 | 0.0257 | 0 | 0.10747 | 0.10747 |
| *Jamaica* | 450 | 0.008046 | 0.030807 | 0 | 0.262963 | 0.262963 |

**Supplemental Table 2, continued**

| Country | Count | Mean Deviation | SD | Min | Max | Range |
|---|---|---|---|---|---|---|
| *Trinidad and Tobago* | 213 | 0.0052 | 0.015503 | 0 | 0.09955 | 0.09955 |
| *South Korea* | 23 | 0.004973 | 0.019298 | 0 | 0.093969 | 0.093969 |
| *Timor-Leste* | 536 | 0.004884 | 0.016336 | 0 | 0.112183 | 0.112183 |
| *Costa Rica* | 692 | 0.004871 | 0.023694 | 0 | 0.213336 | 0.213336 |
| *United Arab Emirates* | 303 | 0.004546 | 0.021288 | 0 | 0.140047 | 0.140047 |
| *Saint Lucia* | 28 | 0.004106 | 0.012973 | 0 | 0.055122 | 0.055122 |
| *Panama* | 1704 | 0.004076 | 0.019316 | 0 | 0.216352 | 0.216352 |
| *Eq. Guinea* | 955 | 0.003257 | 0.017095 | 0 | 0.19235 | 0.19235 |
| *Iran* | 2937 | 0.003238 | 0.020055 | 0 | 0.313523 | 0.313523 |
| *Bahamas* | 425 | 0.002046 | 0.010936 | 0 | 0.104678 | 0.104678 |
| *Vanuatu* | 325 | 0.001953 | 0.014117 | 0 | 0.185129 | 0.185129 |
| *Oman* | 1695 | 0.001612 | 0.012964 | 0 | 0.183969 | 0.183969 |
| *Grenada* | 15 | 0.001142 | 0.002923 | 0 | 0.009292 | 0.009292 |
| *Solomon Is.* | 140 | 0.000664 | 0.005397 | 0 | 0.051616 | 0.051616 |
| *Fiji* | 434 | 0.000282 | 0.002888 | 0 | 0.039222 | 0.039222 |
| *Chile* | 6 | 0 | 0 | 0 | 0 | 0 |
| *Bahrain* | 15 | 0 | 0 | 0 | 0 | 0 |
| *Egypt* | 24 | 0 | 0 | 0 | 0 | 0 |
| *Georgia* | 22 | 0 | 0 | 0 | 0 | 0 |
| *Rwanda* | 7 | 0 | 0 | 0 | 0 | 0 |
| *Montenegro* | 6 | 0 | 0 | 0 | 0 | 0 |
| *Samoa* | 34 | 0 | 0 | 0 | 0 | 0 |
| *São Tomé and Principe* | 30 | 0 | 0 | 0 | 0 | 0 |
| *Niue* | 7 | 0 | 0 | 0 | 0 | 0 |
| *N. Mariana Is.* | 14 | 0 | 0 | 0 | 0 | 0 |
